# Structural and functional evidence for ephaptic control of Purkinje cell spike timing by networks of molecular layer interneurons

**DOI:** 10.64898/2025.12.28.696768

**Authors:** Aliya Norton, Elizabeth P. Lackey, Sevgi Öztürk, Cole S. Gaynor, Sean Ediger, Wei-Chung Allen Lee, Court A. Hull, Wade G. Regehr

## Abstract

Axon collaterals of type 1 molecular layer interneurons (MLI1s) contribute to pinceaux that engulf the initial segments of Purkinje cell (PC) axons and generate extracellular signals that ephaptically inhibit PCs. Here we show that a remarkably large number of MLI1s (∼50) contribute to each pinceau, and that this allows networks of synchronously firing MLI1s to use ephaptic signals to control the precise timing of PC firing *in vivo*.

---

Ephaptic signaling allows neurons to influence each other simply by generating extracellular voltage changes. Even though ephaptic signaling has been observed in numerous brain regions ^1-5^, its role in circuit function remains less well understood than synaptic transmission. *In vitro* studies have suggested that ephaptic signaling may be highly effective at regulating firing for some cell types such as PCs that are sensitive to small extracellular signals because they fire spontaneously and their membrane potential is often near threshold ^6-8^. However, even for PCs it is not known if ephaptic signaling plays a prominent role *in vivo*. Therefore, a deeper understanding of ephaptic signaling onto PCs can both provide insight into how their firing is regulated and establish a model for how ephaptic signaling may regulate activity throughout the brain.

PCs are primarily inhibited by MLIs, which express connexin 36, are electrically coupled, and fire synchronously^9,10^. In addition to making inhibitory GABAergic synapses, MLI1 axons elaborate near the initial segment of PC axons and contribute to specializations known as pinceaux ^10-13^. These pinceaux contain very few vesicles^14,15^ and have low levels of glutamate decarboxylase, bassoon, and the vesicular GABA transporter^14^. In addition, the axon initial segments of PCs lack GABRA1 ^14^, suggesting that pinceaux do not influence PCs by conventional synaptic transmission. In brain slice experiments, paired MLI-PC recordings established that MLIs contributing to a pinceau can ephaptically inhibit connected PCs in the presence of GABA_A_-receptor antagonists ^8,16,17^. However, it is not known if MLI-PC ephaptic signaling is physiologically relevant. In part, this is because it has been difficult to reliably estimate the number of MLI1s that contribute to each pinceau. MLI1s near the bottom of the molecular layer have the classic axonal arborization of basket cells consisting of numerous collaterals that contribute to many pinceaux ^10-13^, MLI1s at intermediate locations in the molecular layer do not look like classic basket cells and contribute to a moderate number of pinceaux ^10,18^, and MLI1s at the top of the molecular layer and type 2 MLIs (MLI2s) do not contribute to pinceaux ^10^. It has been estimated that 3 to 7 MLIs contribute to each pinceau in rodents ^11,19^. However, in the absence of comprehensive serial EM reconstructions such estimates are tentative ^10^. In addition to a lack of information about pinceau connectivity, the properties of MLI1-PC ephaptic signaling have not been tested *in vivo*. Here we find large-scale convergence of MLI1s onto individual PC axon initial segments that can powerfully and selectively regulate PC spike timing, but not overall rate, *in vivo*.

EM reconstructions of a 50 μm-thick volume of lobule V of the cerebellar cortex ^10,20^ were used to estimate the number of MLIs that contribute to each pinceau (**Fig. 1**). Basket cell axons are confined to a 30-μm-thick parasagittal volume ^11^. EM reconstructions are shown for an MLI1 (**Fig. 1a**, *purple*) and a PC with a characteristically extensive dendritic arbor (**Fig. 1a**, *grey*). The cell body of this MLI1 is located near the PC layer and its axon extends parallel to the PC layer and sends a series of collaterals to the bottom of the PC layer that terminate near PC axons. Expanded views show that the MLI1 axon has an elaborate morphology and surrounds the initial segment of a PC axon (**Fig. 1b**) and that glia intercalate the pinceau (**Fig.1c**). The same field of view shows all of the MLI1s that contribute to this pinceau (**Fig. 1d**). Of these MLI1s, 19 were intact, and each of these also contributes to the pinceau of nearby PCs in a parasagittal plane (**Fig. 1e**). There were also 32 large fragments of MLI1 axons that extended to the molecular layer, but these were cut off by the surface of the slice (**Extended data Fig. 1**). An additional reconstructed pinceau along is also provided (**Extended data Fig. 2, Extended data Fig. 3**). On average for each pinceau there are 18.4 ± 1.1 (n=9, range 13 to 22) reconstructed MLI1s that include the soma, 29.5 ± 2.3 (n=9, range 27 to 40) large MLI1 axon fragments, and the sum of complete MLI1s and large fragments is 47.9 ± 2.2 MLI1s (n=9, range 42 to 60) (**Fig. 1f**). A single section with the PC cell body and axon outlined in white (**Fig. 1g**), illustrates the high density of MLI1s near the PC axon initial segment (**Fig. 1hi**). Individual MLI1s are shown in different colors in **Fig. 1h** and are also all shown in purple in **Fig. 1i**. The large fraction of the pinceau comprised of identified MLI1 axons suggests that most MLI1 axons that contribute to the pinceau were detected. MLI1 axons that contribute to a pinceau have complex and highly diverse morphologies, as shown for the images of the MLI1 axons that contribute to the same pinceau from **Fig. 1** (**Fig. 2**). All MLI axons have elaborations near the PC axon, but they range from relatively simple (e.g. **Fig. 2y, mm, ss**) to much more extensive (e.g. **Fig. 2g, l, p**). On some occasions the same MLI1 has two collaterals that contribute to the same pinceau (e.g. **Fig. 2l, r**), raising the possibility that two fragments could arise from the same MLI1. We minimized this issue by restricting our analysis to large fragments that reached the molecular layer. It is likely that excluding small fragments leads to a small underestimate of the number of MLIs per pinceau and having the same MLI1 contribute to 2 different MLI fragments leads to a slight overestimate.

**Figure 1.**
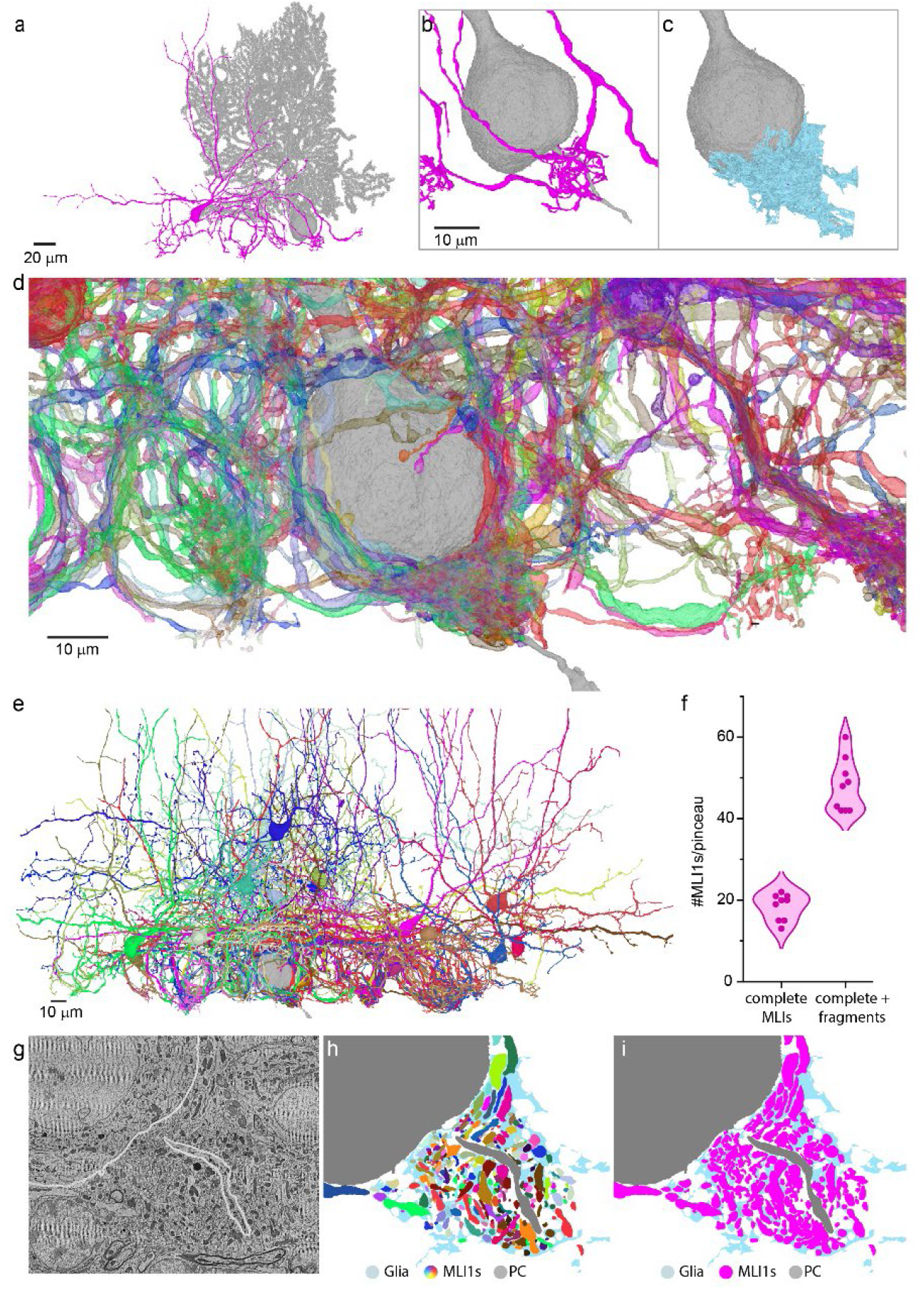
Serial EM reconstructions show that many MLIs contribute to the pinceau onto each Purkinje cell. **a**. Image of an MLI (*purple*) that contributes to the pinceau on the indicated PC (*grey*) **b.** Expanded view of the MLI axon and the PC in **a** **c.** Glia (*light blue*) associated with the pinceau **d.** All MLI axons (n=51) that contribute to the pinceau are shown, with each axon in a different color **e.** Intact MLIs that contribute to the pinceau in **d** are shown (n=19) **f.** Summary of the numbers of MLIs that contribute to the pinceaux associated with 9 different PCs. The number of MLI1s that were reconstructed sufficiently to include the soma (*left*), and the total number of MLI1s including large fragments (**Figure S1**, *right*) are shown **g.** EM section through the center of a pinceau. The cell body and axon of the associated PC are outlined in white **h.** Glia (*light blue*) and MLI1s (*each is a different color*) associated with a pinceau onto the indicated PC (*grey*) **i.** As in **h** but with all MLIs shown in purple.

**Figure 2.**
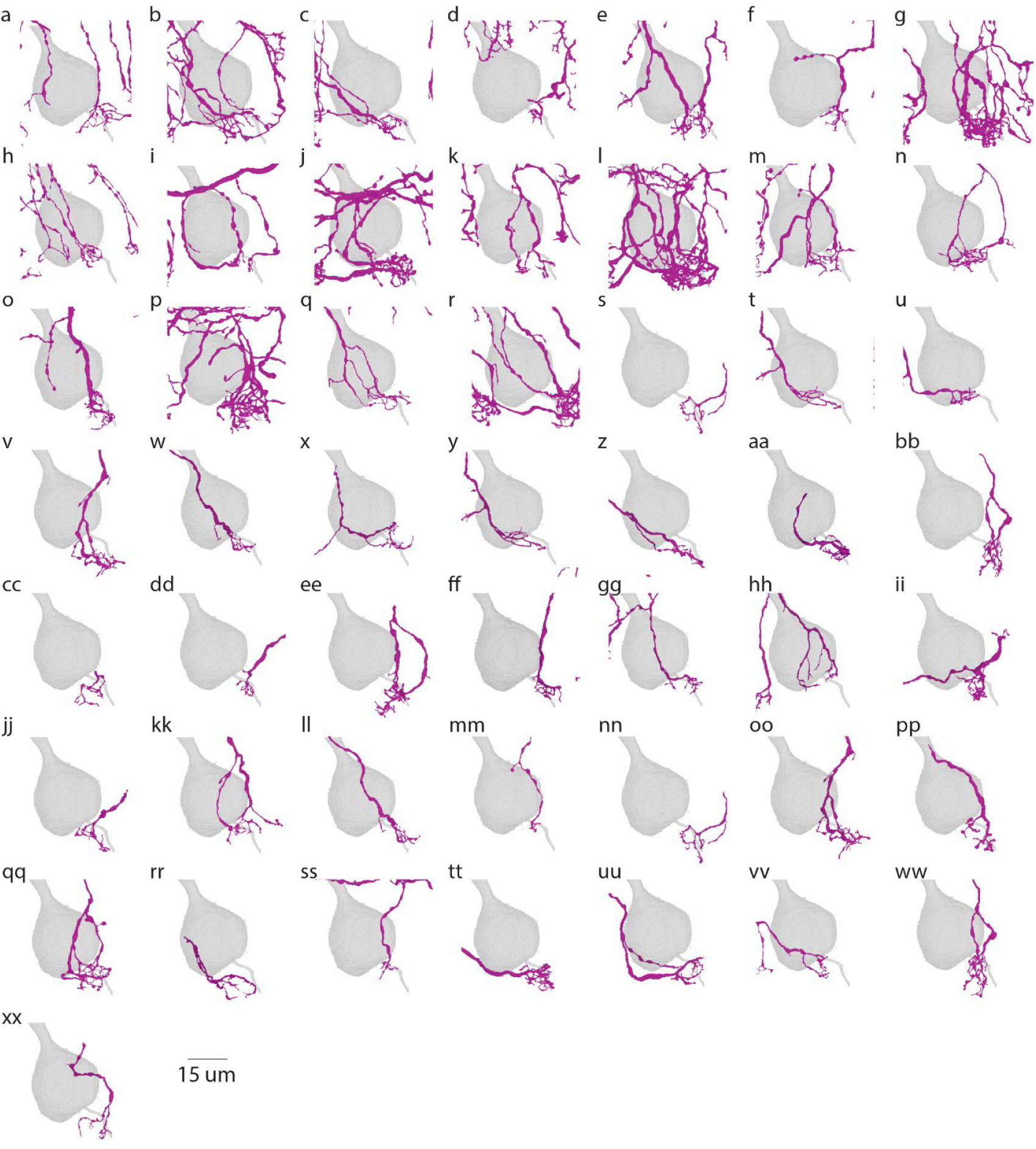
MLIs make diverse contributions to the pinceau of the Purkinje cell in Figure 1 a-xx. Axonal segments of MLIs that contribute to the pinceau in Fig. 1.

To study the functional properties of MLI-PC ephaptic signaling *in vivo* it is necessary to use a different strategy than employed previously in brain slice studies. These used paired MLI-PC recordings with whole-cell recordings from MLIs to both evoke spikes and fluorescently label MLIs, and only observed strong ephaptic suppression of PC firing when the MLI axon was in close proximity to PC axons ^8^. This strategy is impractical *in vivo*, so we first assessed the feasibility of relying on spontaneous firing and MLI1-PC cross correlograms (CCGs) *in vitro*. For cell-attached recordings from spontaneously firing MLIs (**Fig. 3a**, *purple*) and PCs in the presence of GABA_A_ receptor blockers, MLI-PC CCGs show a large and rapid decrease in PC spiking followed by an increase in firing (**Fig. 3a-c**, *blue*), consistent with MLI-PC ephaptic signaling described previously ^8^. The incredible speed of ephaptic signalling was illustrated by the observation that in some cases suppression PC firing began slightly before the detection of a spike in the MLI1 soma, and the increase in firing occurred in less than a millisecond (**Fig. 3d**). Integrating the change in firing revealed that the late increase in firing offset the initial decrease (**Fig. 3e**), and on average there was very little change in the total number of spikes (**Fig. 3f**), with some variability across MLI-PC pairs (**Fig. 3g**) that reflects differences in MLI-PC connections (**Fig. 3h**). These studies suggest that MLI-PC CCGs are suitable for detecting ephaptic signaling *in vivo*.

**Fig. 3.**
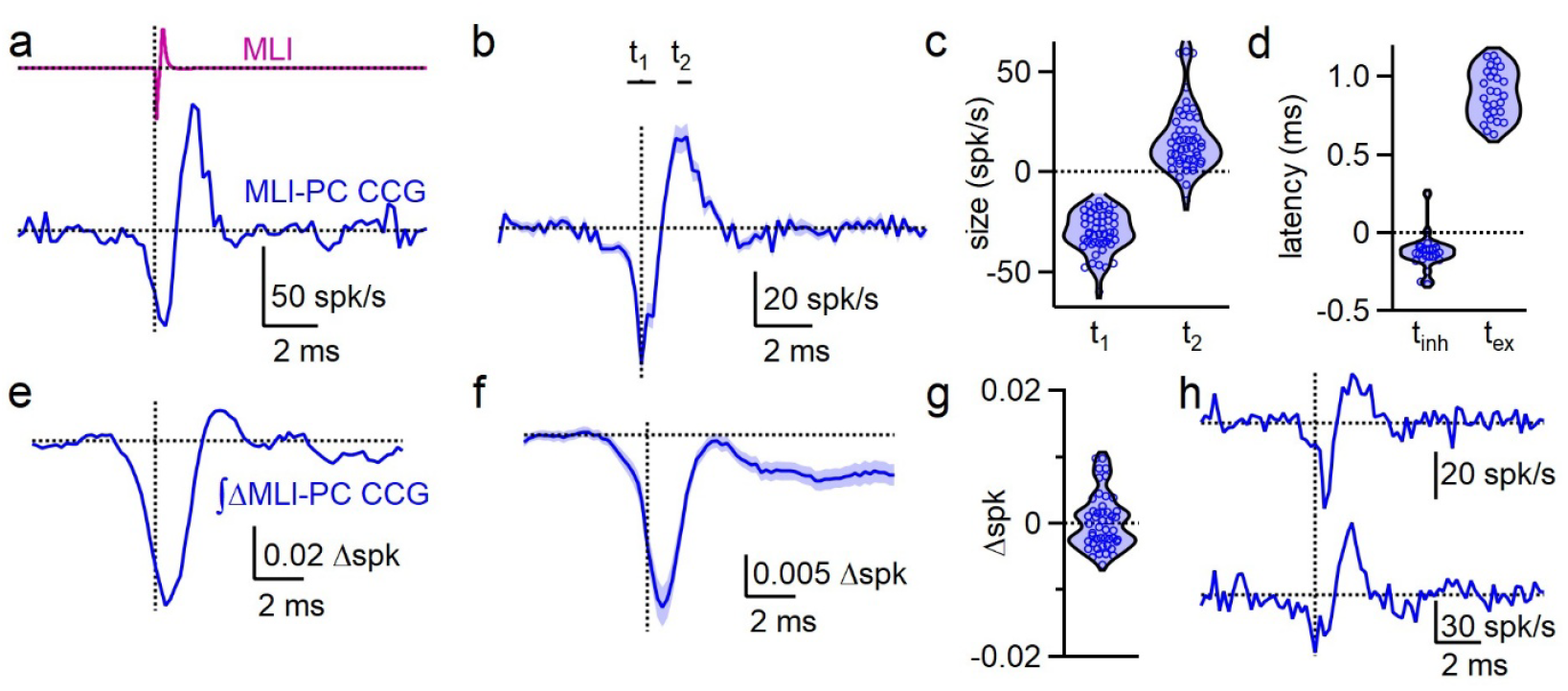
MLI-PC CCGs in spontaneously firing neurons reveal ephaptic inhibition in brain slice. **a.** On-cell recordings from an MLI and a PC were used to determine the MLI-PC cross-correlogram (CCG, *blue*), relative to the MLI spike (*purple*) **b.** Average MLI-PC CCG ± SEM (n=48). t1 and t2 indicate the time intervals summarized in **c** **c.** Summary of amplitudes of components at t1 and t2 for all MLI-PC CCGs **d.** Latencies of the initial suppression (t_inh_) and the subsequent increase (t_ex_) in PC firing (t_inh_: n=24, t_ex_: n=27) **e.** The integral of the MLI-PC CCG was used to assess the net change in PC spiking arising from ephaptic signaling **f.** The average integral of the MLI-PC CCG was used to determine the average net change in PC spiking **g.** The effects of ephaptic signaling on the total number of PC spikes are small and variable **h.** Diverse ephaptic signaling illustrated by example MLI1-PC CCGs.

Neuropixel recordings in awake behaving mice allowed us to determine the properties of ephaptic signaling *in vivo*. In control conditions, most MLI1-PC CCGs consisted of three components: an almost instantaneous suppression, a somewhat delayed suppression, and a late increase in firing (**Fig. 4a**, *red*). Application of AMPAR, NMDAR and GABA_A_R antagonists left the rapid suppression and a later increase in firing (**Fig. 4a**, *blue*) similar to the ephaptic effects observed in brain slice. Subtraction of the CCG in blockers from the initial CCG revealed an inhibitory component (**Fig. 4a**, *black*) that had a longer latency than ephaptic inhibition. Average CCGs were qualitatively similar (**Fig. 4b**). CCGs of individual MLI1-PC pairs showed considerable variability in the amplitudes of ephaptic suppression (**Fig. 4c**), and in later responses (t_2_) that reflect both ephaptic increases in firing and synaptic inhibition (**Fig. 4d**). The latencies of ephaptic and synaptic inhibition were variable, but on average ephaptic inhibition was almost instantaneous, and synaptic inhibition occurred 0.5-1 ms later (**Fig. 4e**). Across pairs, we observed diverse contributions of ephaptic and synaptic components to changes in PC firing (**Fig. 4f,g**). Sometimes the ephaptic and synaptic components are both prominent (**Fig. 4f**, *top*; **Fig. 4g**, *light blue and green*), sometimes the ephaptic component is much larger than the synaptic component (**Fig. 4f**, *middle*; **Fig. 4g**, *red and yellow*), and sometimes the synaptic componenent is much larger than the ephaptic component (**Fig. 4f**, *bottom*; **Fig. 4g**, *dark blue*). On average, MLI1-PC synaptic inhibition decreases the number of PC spikes, but ephaptic suppression affects the timing of PC firing without changing the total number of PC spikes (**Fig. 4h,i**). For individual MLI-PC pairs ephaptic signalling results in small and variable changes in the total number of spikes (**Fig. 4j**), as expected based on MLI-PC CCGs (**Fig. 4k**).

**Figure 4.**
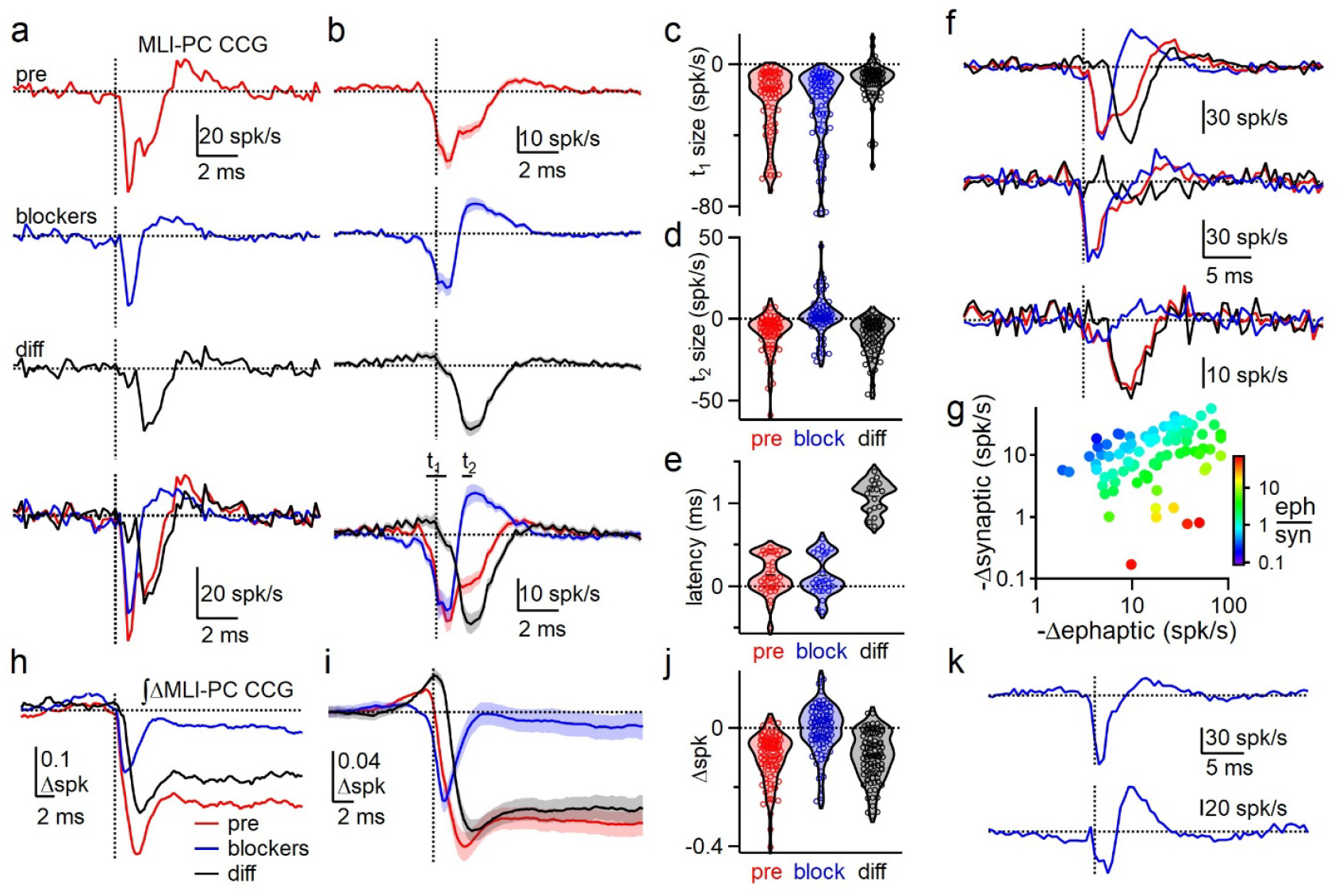
MLI1-PC ephaptic and synaptic inhibition *in vivo*. **a.** An MLI1-PC CCG was measured (*red*) and then AMPARs and GABA_A_ receptors were blocked leaving the ephaptic component (*blue*, NBQX and gabazine). The difference between these components (*black*) was used to estimate the contribution of MLI1-PC synaptic inhibition to the CCG. A superposition of the traces facilitated the comparison of the timing and magnitudes of the different components **b.** Plots as in **a** but for the average of 91 MLI1-PC pairs ± SEM. The two time intervals used for the analysis in **c** and **d** are shown in the lower plot **c.** Violin plots showing the magnitude of the initial component for individual MLI1-PC pairs (n=91) **d.** Violin plots showing the magnitude of the second component for individual MLI1-PC pairs (n=91) **e.** Violin plots showing the timing of the suppression of firing for individual MLI1-PC pairs (pre: n=34, block: n=27, diff: n=17) **f.** Three MLI1-PC CCGs with large ephaptic and synaptic components (*top*), a large ephaptic and small synaptic component (*middle*), and a small ephaptic and large synaptic component (*bottom*) **g.** Scatter plot showing the amplitude of the synaptic inhibition as a function of the magnitude of the initial ephaptic suppression. The synaptic/ephaptic suppression ratios are color coded for each pair **h.** The integral of the change in spiking for the different components is shown for the MLI1-PC pair in a **i.** The integral of the change in spiking for the different components is shown for the average MLI1-PC pair CCG in **b** **j.** Violin plots summarize the effects on spiking for individual MLI1-PC pairs(n=91) **k.** Example MLI1-PC CCGs in the presence of blockers illustrate that ephaptic signaling is dominated by the initial inhibitory component for some pairs *(top*) and by the late excitatory component for others (*bottom*).

These findings fundamentally change the view of MLI1–PC ephaptic signaling. First, they show that approximately 50 MLIs contribute to each pinceau, rather than 3-7 MLIs as was thought to be the case. This suggests that MLI1-PC ephaptic signaling is prevalent and plays an important role in allowing the MLI network to control PC firing. Because MLI1s are electrically coupled and fire synchronously, the large number of pinceax converging on each PC may cooperate to regulate PC firing ^21^. The previous observation that pinceaux are larger in regions with PCs that express zebrin II, and smaller in other region raises the possibility that the number or strength of ephaptic signaling could be regionally specialized ^22^. Second, MLI1-PC CCGs *in vivo* consist of two components: rapid ephaptic inhibition and synaptic inhibition mediated by GABA_A_ receptors that differentially regulate PC firing. MLI-PC ephaptic signaling is very fast and controls PC spike timing without affecting overall spike number, whereas synaptic inhibition is longer-latency and decreases the total number of PC spikes. The finding that MLIs can have such a strong influence on the timing of PC firing *in vivo* raises the possibility that interneurons in other brain regions where axons are in close proximity to the axon initial segments of their targets ^23-26^ might use ephaptic signaling in combination with synaptic transmission to regulate the activity of their targets.

## Online Methods

Serial EM reconstructions were based on a single animal. We previously imaged and aligned a 770 μm X 750 μm X 53 μm volume of lobule V of the mouse cerebellum for EM reconstructions comprised of 1176 45-nm thick parasagittal sections. We used automated image segmentation to generate neuron boundaries^20^.

### Slice experiments

Animal procedures were performed in accordance with the NIH and Animal Care and Use Committee (IACUC) guidelines and protocols approved by the Harvard Medical School Standing Committee on Animals. Animals of either sex were randomly selected for experiments. C57BL/6 mice were obtained from Charles River Laboratories. Animals were housed on a normal light–dark cycle with an ambient temperature of 18–23 °C with 40–60% humidity.

Acute parasagittal slices (220-μm thick) were prepared from P28-45 mice. Mice were anaesthetized with an intraperitoneal injection of 100 mg/kg ketamine + 10 mg/kg xylazine and perfused transcardially with an ice-cold solution containing (in mM): 110 choline chloride, 7 MgCl_2_, 2.5 KCl, 1.25 NaH_2_PO_4_, 0.5 CaCl_2_, 25 glucose, 11.6 sodium ascorbate, 3.1 sodium pyruvate, 25 NaHCO_3_, equilibrated with 95% O_2_ and 5% CO_2_. Slices were cut in the same solution and then transferred to artificial cerebrospinal fluid (ACSF) containing (in mM) 125 NaCl, 26 NaHCO_3_, 1.25 NaH_2_PO_4_, 2.5 KCl, 1 MgCl_2_, 1.5 CaCl_2_ and 25 glucose equilibrated with 95% O_2_ and 5% CO_2_. Following incubation at 34 °C for 30 min, the slices were kept up to 6 h at room temperature until recording.

Recordings were performed at 32 °C. Visually guided on-cell recordings were obtained with patch pipettes of 1-2-MΩ resistance for PCs and 3-5-MΩ resistance for MLIs pulled from borosilicate capillary glass (BF150-86-10, Sutter Instrument). Glass pipettes were filled with ACSF for on-cell recordings. Electrophysiology data were acquired using a Multiclamp 700A or 700B amplifier (Axon Instruments), digitized at 100 kHz and filtered at 10 kHz. Acquisition and analysis of slice electrophysiological data were performed using custom routines written in Igor Pro (Wavemetrics) and Matlab. The following receptor antagonists were added to the ACSF solution to block glutamatergic, GABAergic, and glycinergic synaptic currents (in μM): 2.5 (R)-CPP, 5 NBQX, 10 gabazine, 1.5 CGP, 1 strychnine. All drugs were purchased from Abcam and Tocris.

Recordings were made from lobules IV-V of the vermis. Spontaneous action potentials from MLI-MLI-PC triplets were recorded in loose-patch configuration for 5-10 min. MLI pairs were located within 5 µm of each other in the sagittal plane and within 100 µm of the PC. MLI position was measured from the bottom of the PC layer. Spikes were detected in IGOR Pro and manually verified for each cell. Spike times were aligned to the peak of the first derivative of the action potential. Cross-correlograms were subsequently calculated from the binarized spike trains using a 0.2 ms bin size. This is equivalent to counting the number of spikes in the PC which fall in bins at different time lags around the spikes of the MLI. Bursting PCs (5/80 MLI-PC pairs) were excluded from the analysis.

For MLI-PC cross-correlograms from slice recordings, spike suppression was considered significant if the z-score crossed <-3. For connected pairs, the amplitude of MLI-PC ephaptic inhibition was measured as the minimum PC Δspk/s from t = -0.5 to 0.5 ms (t_1_), and the magnitude of excitation was measured as the average over a 0.5 ms window centered at 1.5 ms (t_2_). All responses were measured relative to baseline averaged 20 ms prior to the MLI spike. Latency was measured as the half max time for t_1_ and t_2_ for response amplitudes with z-scores <-5 or >5 for each component. The cumulative changes in PC spikes were measured as the average of the integrals of the cross-correlograms from 3-5 ms following the MLI spike.

### In vivo experiments

Mice for in vivo recordings (5 C57xCBA mice, 3 male, > P85) were housed in a normal (12 hr light/ 12 hr dark) cycle. All *in vivo* experimental procedures were performed with approval from the Duke University Animal Care and Use Committee.

### Surgical Procedures

Mice underwent headpost implantation surgery for head-fixed preparation prior to electrophysiological recordings. An initial dose of ketamine/xylazine (50 mg/kg and 5 mg/kg IP) and meloxicam (5 mg/kg subq) were injected 20 min before the induction of isoflurane. Isoflurane (1-2 %) was administered throughout the surgery to maintain deep anesthesia, with anesthetic depth verified every 30 min by the absence of a paw-pinch withdrawal reflex. Body temperature was maintained at 35-37 degree Celsius using a heating blanket (TC-111 CWE). Eyes were protected with ophthalmic ointment (GenTeal). Animals were secured in a stereotaxic frame using ear bars, and the scalp was shaved with depilatory cream (Nair) and cleaned with betadine and 70% ethanol. After a midline incision, the scalp was retracted, and the skull was cleaned, dried, and gently scraped. A titanium headpost (HE Palmer) was placed onto the skull together with a stainless steel ground screw (F.S. Tools) over the left cerebellum and affixed with Metabond (Parkell). Buprenorphine (0.05 mg/kg SQ) was administered every 12 hours for 48 hours after surgery and mice were monitored for 4 days.

### In vivo electrophysiology

Mice were habituated for head-fixation on the wheel after +7 days of headpost surgery. Mice were given dexamethasone (3 mg/kg SQ) 3-12 hours before each recording session. Craniotomies (∼ 1 mm) were drilled under 1-1.5 % isoflurane anesthesia over lobule simplex ( 1-2 mm lateral; 6-6.5 mm posterior to Bregma). Additional craniotomies adjacent to the initial site were drilled as needed for subsequent recordings and were sealed between sessions using Qwik-Cast (WPI) and Metabond. After craniotomy was prepared, mice were head-fixed on the wheel and allowed to recover from anesthesia for at least 2 h before recordings began. Neuropixels 1.0 electrodes were lowered through the cerebellar cortex at a rate of 1 μm/sec, advanced ∼250 μm beyond the target depth, and then retracted by ∼250 μm to relieve tissue tension. Probes were allowed to stabilize for at least 20 min with ASCF applied over the craniotomy prior to recordings. Units recorded between 0 and -2200 μm were included in the analyses. Wheel position was monitored using a rotary encoder (YUMO).

Recordings consisted of a 30-min baseline period, followed by 20-min drug wash-in period and 30-40 min of recording during synaptic block. After the baseline recording, ASCF was removed from the craniotomy and a cocktail of synaptic blockers (300 μM gabazine, 800 μM NBQX and 800 μm AP5) was applied. On the final recording day, the non-recording side of the probe was coated with dye (DiI) to enable post hoc verification of the electrode location. Animals were then deeply anesthetized with ketamine/xylazine (300 mg/kg and 30 mg/kg, IP) and perfused for histology.

Recordings were made using data acquisition system consisting of National Instruments hardware (NI PXIe-1071 for neural data and NI SCB-68A for movement signals) and SpikeGLX software (v.20200520 from billkarsh.github.io).

Post-processing included demultiplexing (CatGT 3.5), alignment of behavioral timepoints to neural data timepoints (TPrime 1.8), spike sorting (Kilosort 4.0), high-pass filtering of raw voltage traces at 300 Hz using NeuroPyxels v.2.0.4 (Beau et al., 2021, Zenodo), cell-type classification using C4 classifier (Beau et al., 2025 https://www.c4-database.com/apps/classifier), and manual curation with Phyllum (GitHub -blinklab/phyllum: a custom phy plug-in for cerebellar layer identification developed in the Medina lab at Baylor College of Medicine). Units were selected for further analyses only if they exhibited <5% refractory period violations and maintained stable amplitude distributions throughout the recording.

### MLI subtype identification

MLI1s were identified following the criteria described previously (Lackey et al. 2024). In the present work, we additionally incorporated the C4 classifier (Beau et al., 2025), a deep-learning-based algorithm that identifies cerebellar cell types using laminar location, waveform features, and 3D auto-correlograms (ACGs). After MLIs were identified using the C4 classifier, we further subclassified MLI1s based on their interactions with PCs and other MLIs according to Lackey et al. MLIs that fired > 4Hz, produced strong short-latency inhibition of a PC (>= 3 STD, latency <4 ms), and did not inhibit other MLIs were classified as MLI1s. In total, 13 recordings yielded 100 MLIs, of which 26 were classified as MLI1s.

### PC-MLI1 cross-correlograms

For MLI-PC cross-correlograms from in vivo recordings, spike suppression was considered significant if the z-score crossed <-3. For connected pairs, the changes in PC spike rates following the MLI spike were measured as the minimum from t = -0.5 to 0.5 ms (t_1_), and the average over a 0.5 ms window centered at t = 1.5 ms (t_2_). The cumulative changes in PC spikes were measured as the average of the integrals of the correlograms from 10-15 ms following the MLI spike. All responses were measured relative to baseline averaged 20 ms prior to the MLI spike. Latency was measured as the half max time for amplitudes with z-scores <-5 or >5 for each component.

## Data and Code Availability

Datasets supporting the findings of this study are available from the corresponding author upon request. Custom code used in this work will be posted on GitHub. Any additional information is available from the corresponding authors upon request.

## Acknowledgments

This work was supported by the NIH (R35NS097284 and R35142971 to W.G.R., F32NS133036 to E.P.L., R21NS085320, RF1MH128949, and RF1MH114047 to W.-C.A.L., R01NS128054 and R01NS141997 to C.A.H.), the Nancy Lurie Marks Foundation (to E.P.L.), the Ruth K. Broad Biomedical Research Foundation (388000038 to M.E.H.), the Bertarelli Program in Translational Neuroscience and Neuroengineering, Stanley and Theodora Feldberg Fund, and the Edward R. and Anne G. Lefler Center. Portions of this research were conducted on the O2 High Performance Compute Cluster at Harvard Medical School.

