## Supplemental Figures for "Structural and functional evidence for ephaptic control of Purkinje cell spike timing by networks of molecular layer interneurons"

**Supplementary Figures**

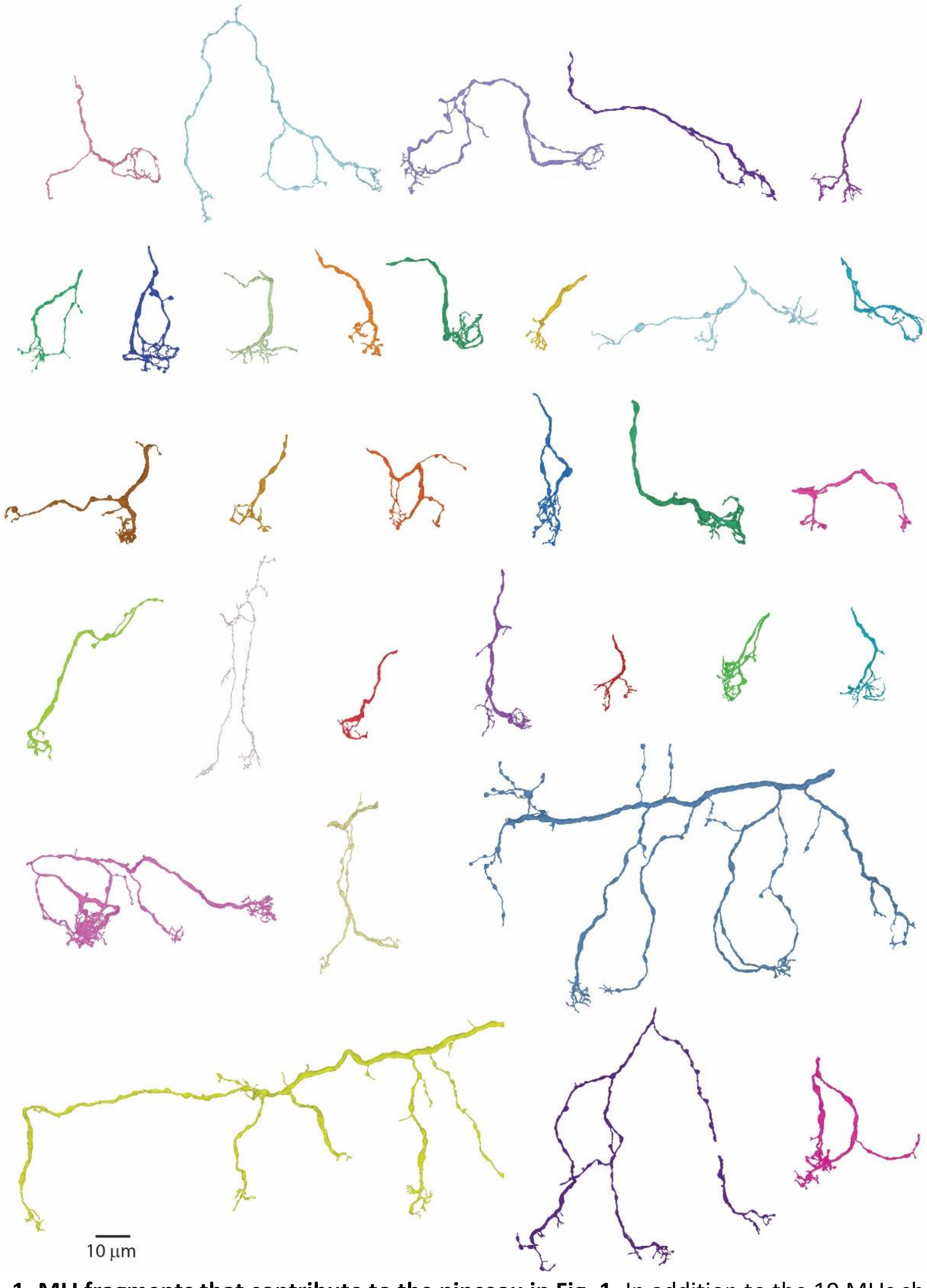


**Extended data Fig. 1. MLI fragments that contribute to the pinceau in Fig. 1.** In addition to the 19 MLIs shown in **Fig. 1e,** these 32 fragments contributed to the pinceau shown in **Fig. 1d**.

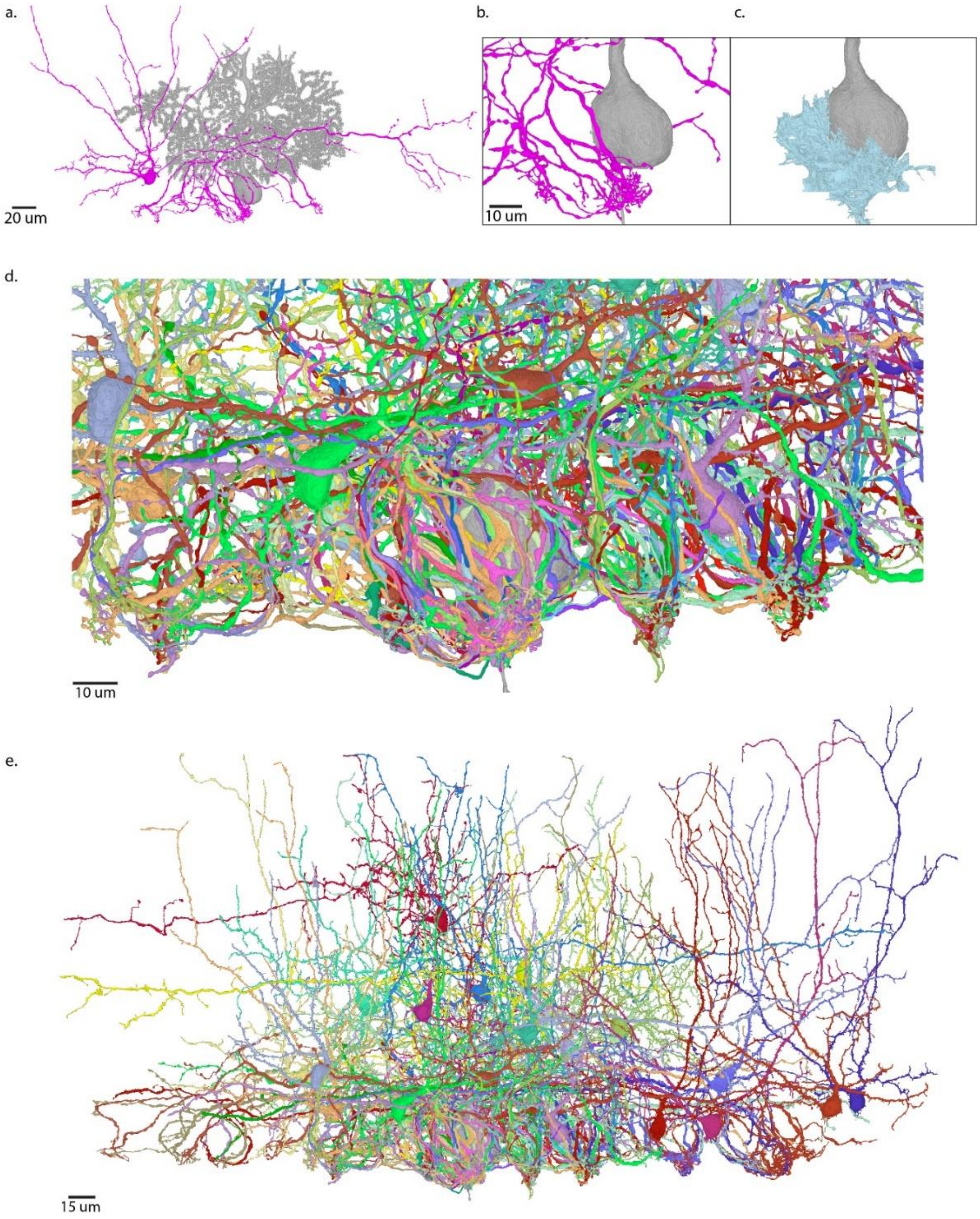


**Extended data Fig. 2. Another serial EM reconstruction showing MLIs that contribute to the pinceau onto the PC.**
**a.** Image of an MLI (*purple*) that contributes to the pinceau on the indicated PC (*grey*).
**b.** Expanded view of the MLI axon and the PC in **a**.
**c.** Glia (*light blue*) associated with the pinceau.
**d.** All MLI axons that contribute to the pinceau are shown, with each axon in a different color.
**e.** Intact MLIs that contribute to the pinceau in **d** are shown.

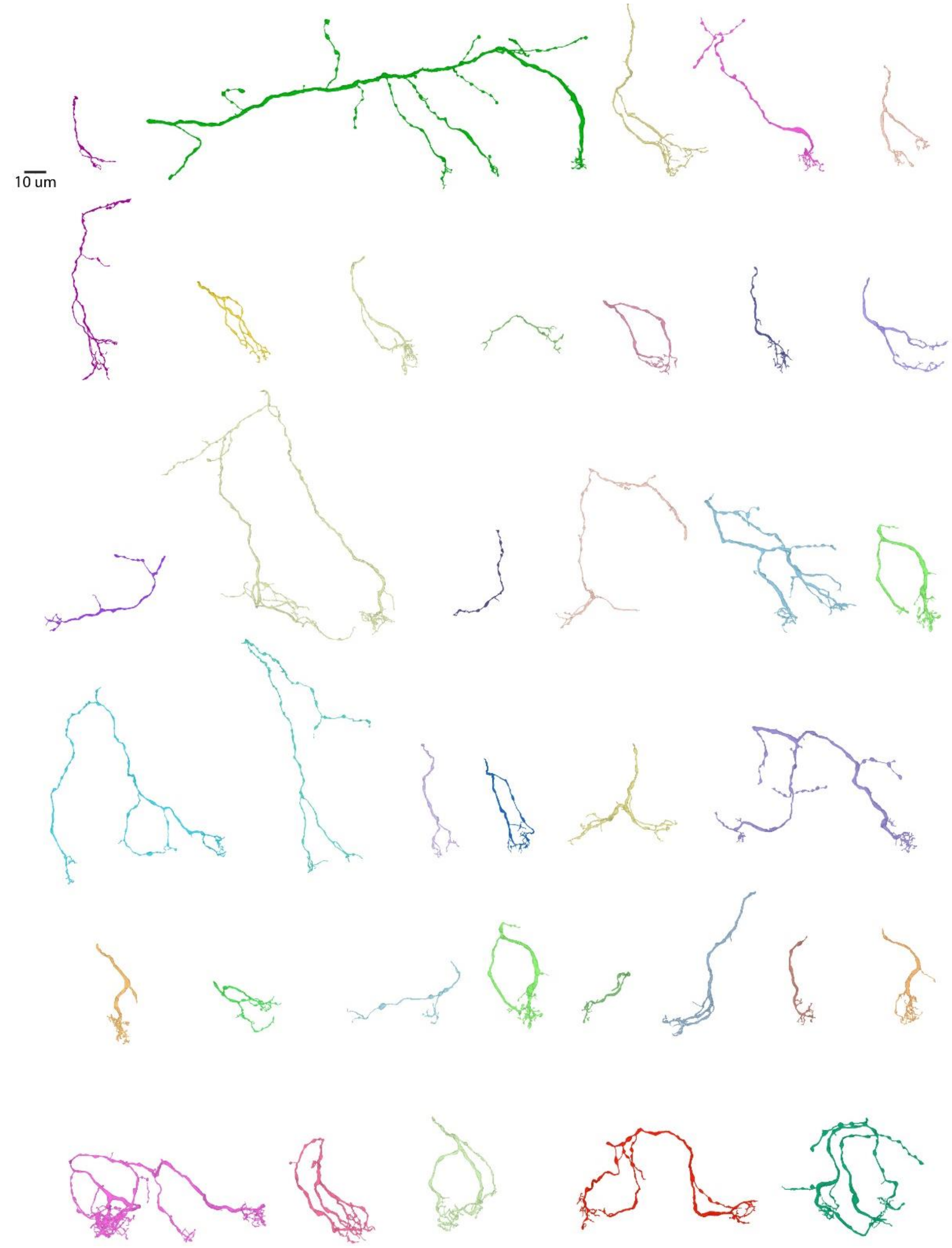


**Extended data Fig. 3. MLI fragments that contribute to the pinceau in Extended data Fig. 2.** In addition to the MLIs shown in **Fig. S2e**, these 38 fragments contributed to the pinceau. Note that some of the fragments are the same as in Extended data Fig. 3 because these two pinceaux share MLI1 inputs.
